# Tailocin-mediated interactions among Soft Rot *Pectobacteriaceae*

**DOI:** 10.1101/2024.09.28.615580

**Authors:** Marcin Borowicz, Dorota M. Krzyżanowska, Marta Sobolewska, Magdalena Narajczyk, Inez Mruk, Paulina Czaplewska, Jacques Pédron, Marie-Anne Barny, Pierre Yves Canto, Joanna Dziadkowiec, Robert Czajkowski

## Abstract

Bacteria carry phage-derived elements within their genomes, some of which can produce phage-like particles (tailocins) used as weapons to kill kin strains in response to environmental conditions. This study investigates the production and activity of tailocins by plant pathogenic bacteria: *Pectobacterium*, *Dickeya,* and *Musicola* genera, which compete for niche, providing an attractive model to study the ecological role of tailocins. Microscopy revealed that most analyzed strains (88%) produced tailocins. Tailocin-mediated killing interactions were assessed across 351 strain pairs, showing that *Dickeya* spp. had a higher likelihood of killing neighbors (57.1%) than *Pectobacterium* spp. (21.6%). Additionally, *Dickeya* spp. strains exhibited broader phylogenetic killing, targeting both *Pectobacterium* spp. and *Musicola* sp., while *Pectobacterium* spp. tailocins were genus-specific. Mutual killing was observed in 33.9% of interactions, predominantly within *Dickeya* spp. Although tailocins were morphologically indistinguishable between producers, genomic analyses identified conserved clusters having distinct differences between *Pectobacterium* spp. and *Dickeya* spp tailocins. This suggests different origins of these particles. Induction experiments demonstrated that tailocin production was boosted by hydrogen peroxide, supporting the role of these particles in bacteria-bacteria competition during infection. Tailocins were detectable in infected potato tissue but not in river water, highlighting the particular ecological relevance of tailocins in plant environments.

## Introduction

In highly competitive and resource-limited environments that bacteria occupy, their survival and fitness often depend on their ability to outcompete neighboring rival cells (Bauer et al., 2018;Wagner, 2022). To gain a competitive edge, bacteria have evolved diverse strategies, including producing and utilizing phage tail-like particles (tailocins) that can be lethal to other bacteria. These structures directly resemble the tails of bacteriophages but function independently from viral infection and replication (Booth et al., 2023). These particles are highly specialized killing instruments that, by puncturing the cell envelope, lead to the disruption of the cell membrane and the death of the targeted cell (Scholl, 2017). Until recently, the utilization of phage tail-like particles has been primarily studied in *Pseudomonas* species (Carim et al., 2021). However, there are also reports of tailocins isolated from various other Gram-negative and Gram-positive bacterial species, including human, animal, and plant pathogens, as well as saprophytic bacteria that inhabit a wide range of environments (Backman et al., 2024a).

We previously characterized a new type of phage tail-like particle produced by the plant pathogenic bacterium, *Dickeya dadantii* reference strain 3937 (Borowicz et al., 2023). This tailocin, dickeyocin P2D1, was tolerant of most environmental conditions and exhibited broad killing potential against members of different *Dickeya* species but was harmless to *Pectobacterium* species and non-toxic to *Caenorhabditis elegans.* We postulated that the dickeyocin P2D1 targets were restricted to phylogenetically closely related strains. Only one other phage tail-like particle, carotovoricin Er from *Pectobacterium carotovorum* Er, was described in detail (Yamada et al., 2006). The sequence of Er differs from that of dickeyocin P2D1, indicating a distinct evolutionary origin of these two tailocins (Yamada et al., 2006). Besides these limited studies, little attention has been paid to the presence and ecological role of tailocins in SRP bacteria.

SRP bacteria (consisting of *Pectobacterium* spp., *Dickeya* spp., and *Musicola* spp., previously collectively described as pectinolytic *Erwinia* spp.) are a particularly appropriate group of bacteria for studying the environmental role of phage tail-like particles in the environment. *Pectobacterium* spp., *Dickeya* spp., and *Musicola* spp. belong to the same species complex and are often found together in infected hosts. Consequently, due to their metabolic similarities and overlapping niches, they are known to compete against each other (Toth et al., 2021a). Furthermore, these bacteria occupy various ecological niches, including host and non-host plants, natural and agricultural soils, rainwater, surface water, sewage, and insects (Van Gijsegem et al., 2021). In these diverse environments, SRP bacteria often encounter a diverse array of other species, both closely and distantly related, but their competitive strategies in these alternative biomes remain unknown (Bellieny-Rabelo et al., 2019).

Our understanding of the mechanisms of competition in SRP bacteria mediated by phage tail-like particles remains poorly understood (Marquez-Villavicencio et al., 2011;Hugouvieux-Cotte-Pattat, 2016;Hugouvieux-Cotte-Pattat et al., 2023). Specifically, no systematic studies have been conducted to assess whether the production of tailocins is common in this group of bacteria and how it contributes to their overall ecological success (Borowicz et al., 2023). Addressing this gap remains important for understanding how plant-pathogenic bacteria establish and thrive in diverse environments, particularly in those niches where microbial competition between kin strains is expected to be intense.

This study aimed to assess the expression of phage tail-like particles (tailocins) in SRP strains obtained from one selected environment – the Durance River in France (Ben Moussa et al., 2022). Although river water is not recognized as a primary environment where SRP bacteria thrive, this niche links the several distinct ecosystems that support the growth and survival of SRP. The tested SRP strains could potentially interact in water or be transmitted to a common plant host, where they might interact during infection (Van Gijsegem et al., 2021). We aimed to characterize interactions among SRP bacteria guided by phage tail-like particles to understand better how these pathogens share the environment and whether they compete for niche during infection. For this, we characterized the SRP tailocins and identified gene clusters associated with their synthesis, thus determining the prevalence of tailocin production among environmental SRP strains and their apparent role in intragenus competition among the diverse strains that coexist in this aquatic habitat.

## Materials and Methods

### Bacterial strains and culture conditions

Bacterial strains included in this work are listed in Supplementary Table S1. The pool comprised 25 SRP isolates from the waters of Durance River (France), collected between 2015 and 2017 (Ben Moussa et al., 2022), as well as two reference strains: *Dickeya dadantii* 3937 (Pritchard et al., 2013) and *Musicola paradisiaca* NCPPB 2511 (Hugouvieux-Cotte-Pattat et al., 2021). The latter strains have been previously studied (Borowicz et al., 2023). Routine propagation of bacterial cells was performed at 28 °C in liquid Trypticase Soy Broth (TSB, Oxoid), with agitation (120 rpm), or on solid Trypticase Soy Agar (TSA, Oxoid) under the same growth conditions (Czajkowski et al., 2010).

### Induction and purification of tailocins

Tailocin particles were purified as described previously (Borowicz et al., 2023). Briefly, an overnight culture (*ca.* 16 h) of each strain was rejuvenated (1:40) in a fresh aliquot of TSB and cultured for 2.5 hours. Next, tailocin production was induced by the addition of 0.1 μg mL^−1^ of mitomycin C (Abcam, Poland). Twenty-four hours after induction, tailocins were purified using PEG from 10 mL volumes of filtered (0.2 µm, PES membrane, GoogLab™) culture supernatants. Purified particles were stored in phosphate buffer saline (PBS), pH 7.2, at 4 °C.

### Imaging

The morphology of purified tailocin particles was investigated by transmission electron microscopy (TEM) and atomic force microscopy (AFM). TEM imaging was performed as described previously (Borowicz et al., 2023) using a Tecnai Spirit BioTWIN microscope (FEI). AFM imaging was performed using JPK NanoWizard® 4 (NanoScience), in non-destructive quantitative imaging (QI™) force spectroscopy mode, employing the SCANASYST-AIR probes (f0 7.0 kHz, diameter <12 nm, k: 0.4 N/m) and the SCANASYST-FLUID+ probes (f0 150 kHz, diameter <12 nm, k: 0.7 N/m). Spring constants and sensitivity of the probes were calibrated using the thermal calibration method. Before imaging, the samples were deposited on freshly cleaved mica surfaces and air-dried. Images were processed using JPK Data Processing software. Dimensions of tailocins were calculated based on TEM images. Tailocins from a given strain were considered for measurement only if more than 4 individual particle images were available for analysis. Analysis was accomplished on 13 out of 18 *Pectobacterium* spp. strains, with a median of 10 particles measured per strain, and 7 *Dickeya* spp., with a median of 17 particles measured per strain.

### Killing and sensitivity assays

Twenty-seven SRP strains, including 25 water isolates (Supplementary Table S1), were tested in pairs to determine the target range (killing spectrum) of the tailocin produced by each group member. Likewise, the sensitivity of strains to the tailocins produced by all other strains in the analyzed pool was assessed. Both the killing spectrum and sensitivity were tested using the spot assay as previously described (Hockett and Baltrus, 2017;Yao et al., 2017;Borowicz et al., 2023). Each combination was tested in duplicate, and the entire experiment was performed twice using independently-obtained batches of tailocins.

### SDS-PAGE, ESI LC-MS/MS, and mapping of identified proteins to genomic *loci*

Proteins in PEG-purified tailocin preparations were separated by SDS-PAGE, fragmented, and the resulting peptides sequenced using mass spectrometry, as described in detail previously (Golebiowski et al., 2022;Borowicz et al., 2023). Briefly, bands were excised from the gel and subjected to in-gel trypsin digestion, and the peptides were eluted, cleaned up, and concentrated (Schmidt and Sinz, 2017;Goldman et al., 2019). The peptides were then analyzed by ESI LC-MS/MS. This included separating peptides on an Eksigent microLC column ChromXP C18CL (3 μm, 120 Å, 150 × 0.3 mm). The samples were placed onto the column with a CTC Pal Autosampler (CTC Analytics AG, Zwinger, Switzerland), and the injection volume was 5 μl. Solvents A and B consisted of 0.1% (v/v) formic acid (FA) in water and acetonitrile, respectively. LC gradient parameters: 18–50% solvent B in 30 min (Fiołka et al., 2019). The mass spectra with Triple Tof 5600+ mass spectrometer with DuoSpray Ion Source (AB SCIEX, Framingham, MA, USA) enabled the identification of proteins based on the obtained fragmentation spectra using the Peaks Studio 11 (Bioinformatic Solutions) and the appropriate protein database (Supplementary Table S1) (1% FDR). The annotations of the identified proteins were searched for phage-associated functions, and the encoding genes were mapped to the genomes of the tested strains. Tailocin-encoding regions were identified based on the clustering of the identified phage/tailocin-related genes within the genomes of the analyzed strains and previous studies concerning the structure of the P2D1 tailocin cluster (Borowicz et al., 2023). As the full annotated sequence of the carotovoricin Er cluster is unavailable, the originally uploaded sequence (accession: AB017338.2) (Yamada et al., 2006) annotated by us using RAST (https://rast.nmpdr.org/rast.cgi) (Brettin et al., 2015) was used for cluster synteny analysis.

### Phylogenetic and bioinformatic analyses

#### Synteny of tailocin clusters

Gene clusters encoding tailocins were extracted from the genomes of *Pectobacterium*, *Dickeya,* and *Musicola* species except for *Pectobacterium odoriferum* A122-S21-F16, where the putative cluster was split across multiple contigs. For *Pectobacterium* spp. the genomic region containing the tailocin cluster was defined as the region between the genes annotated as *tolC* (locus tag in *P. versatile* A69-S13-O15: EG331_00440) and *ybiB* (locus tag in *P. versatile* A69-S13-O15: EG331_00565). In contrast, for *Dickeya* species and a single *Musicola* sp. strain, the genomic region containing the tailocin cluster was identified as the region between the genes annotated as *methyl-accepting chemotaxis protein* (locus tag in *D. dadantii* 3937: DDA3937_RS1200) and *guaD* (locus tag in *D. dadantii* 3937: DDA3937_RS12145).

Accession numbers of genomes from which the tailocin regions were extracted are listed in Supplementary Table 1. The extracted sequences were then subjected to synteny analysis and visualization using the Clinker tool (Gilchrist and Chooi, 2021). The analysis was performed separately for *Pectobacterium* and *Dickeya* species, along with a single strain of *Musicola* sp. Genes within the clusters were color-coded based on their putative roles in tailocin production. In the visualization, links between genes indicate sequence identity percentages greater than 50%. The order of strains in the visualization was determined according to their phylogenetic relationships. Synteny analysis was based on *Pectobacterium versatile* A73-S18-O15 for *Pectobacterium* spp. and *Dickeya dadantii* 3937 for *Dickeya* spp.

#### Multilocus sequence analysis

A subset of strains was used for phylogenetic analysis. The selected strains possessed the following features: (1) the tailocin cluster had to be present on a single genomic contig; (2) genes annotated as fiber and sheath had to be present within the region identified as the tailocin cluster; and (3) the strain had to demonstrate the ability to target at least one other analyzed strain through tailocin interaction. Orthologous sequences were clustered into homologous families using the SiLix software package v1.2.9 (Miele et al., 2011) with an 80% identity threshold and at least 80% overlap. Orthologous sequences (658 coding sequences) were concatenated, aligned using MUSCLE (Edgar, 2004) software v5.1, and filtered using the GBLOCK tool (Castresana, 2000). The alignments were used to build a phylogenetic tree with the BioNJ algorithm with SeaView software v5.0.5 (Gouy et al., 2010), with 200 bootstrap replications.

#### Phenotype-based dendrogram

Agglomerative clustering (Nielsen, 2016) was applied to prepare a dendrogram for tested strains based on the similarity of their killing spectra. The ability to kill a given strain was assigned the value of 1, and the lack of this ability was noted as 0. The scikit-learn (Virtanen et al., 2020) and SciPy (Virtanen et al., 2020) libraries were used for clustering and dendrogram plotting, respectively. A Phyton script for implementing the above libraries was written with the assistance of ChatGPT-4o (OpenAI) (Supplementary Dataset 3).

#### *Prevalence of carotovoricin and dickeyocin clusters in* SRP complete genomes

The carotovorocin cluster was blasted against 116 *Pectobacterium* spp. complete, high-quality genomes available on the NCBI database in June 2024 using the NCBI Megablast algorithm with default parameters (https://blast.ncbi.nlm.nih.gov/Blast.cgi). The presence of the cluster was defined according to the 70% sequence coverage and identity thresholds. The available sequence of the originally described carotovoricin Er cluster was incomplete (accession: AB017338.2). Therefore, as a reference carotovoricin sequence for *Pectobacterium* spp. we used the full nucleotide sequence of the carotovoricin cluster from *P. versatile* A73-S18-O15 (19,528 bp.). For *Dickeya* spp. and the single *Musicola* sp. strain, the full nucleotide sequence of the dickeyocin cluster of *D. dadantii* 3937 strain (26,580 bp) was blasted against a single *Musicola* sp. and 74 *Dickeya* spp. complete genomes available on the NCBI database in June 2024. Results were interpreted based on analogous thresholds as described for P2D1 tailocin.

### Detection of tailocins in river water and potato tubers

To determine whether SRPs produce detectable levels of tailocins in river water and potato tuber tissue, we inoculated these environments with bacteria. After incubation, the tailocins were purified and spot tests were conducted as outlined below. Each experiment was performed twice, and detection was considered positive when all replicates gave positive results. To prepare the inoculation suspensions, the six strains *P. versatile* A69-S13-O15, *P. quasiaquaticum* A411-S4-F17, *P. aquaticum* A212-S19-A16, *D. dadantii* 3937, *D. chrysanthemi* A604-S21-A17, and *D. zeae* A586-S18-A17, were grown in TSB (Oxoid) for 24 h at 28 °C. The cells were harvested by centrifugation (4200 RCF, 5 min) and resuspended in PBS buffer. Turbidity of the suspension was adjusted to 0.06 McFarland (approx. 10^6^ CFU mL^−1^). Five ml of filter-sterilized (0.2 µm, PES membrane) and autoclaved Durance River water was inoculated with 50 µl of bacterial suspension, three replicates per strain. Uninoculated water was used as a control. Samples were incubated for 72 h at 15 °C with shaking (120 rpm) to mimic natural conditions. Following incubation, 2 ml of water from each sample was centrifuged to remove bacterial cells (8000 RCF, 10 min), and the supernatant was processed for purification and detection of tailocins. Tubers cv. Gala were surface sterilized by immersion for 20 min in 5% commercial bleach (ACE, Procter and Gamble), followed by a double rinse in distilled water and air drying under laminar flow. Each tuber was inoculated by inserting a pipette tip containing 50 µl of the test suspension (Krzyzanowska et al., 2019). Tubers inoculated with PBS buffer alone were used as a negative control. Three potato tubers were tested per treatment. Inoculated tubers were placed in humid boxes (85 to 90% relative humidity). Samples were incubated at 28°C for 72 h to enable the development of soft rot symptoms. The tubers were then cut at the inoculation site. Tissue macerated by bacteria was excised, weighed, and placed in homogenization bags (Bioreba). PBS buffer supplemented with 0.02% diethyldithiocarbamic acid (DIECA) (Sigma-Aldrich) was added to the samples (1:2, w/v) (Perombelon and van Der Wolf, 2002). The samples were homogenized, and 1 mL aliquots were diluted 1:1 with PBS 0.02% DIECA in new tubes. Bacteria and plant debris were pelleted by centrifugation (8000 RCF, 10 min, RT), and the supernatant was processed for purification and detection of tailocins. The supernatants from both experimental setups were filtered (0.2 µm, PES membrane) and transferred to new tubes containing PEG-8000 (final concentration 10 % m/v). Samples were incubated overnight at 4°C with shaking. The putative tailocins were collected by centrifugation (1 h, 16 000 RCF, 4°C) and resuspended in 1/20 volume of the initial sample in PBS. Tailocins were detected using a spot test (Hockett and Baltrus, 2017;Yao et al., 2017). Five µL aliquots of purified tailocins were spotted on soft top agar plates, each inoculated with a strain sensitive to tailocins of the six investigated SRPs (Supplementary Table S2).

### Semiquantitative assessment of tailocin production depending on the type of inductive factor

We investigated whether hydrogen peroxide can induce the production of tailocins in SRPs in a way similar to mitomycin C. The experiment was conducted on the same group of six strains tested for tailocin production in river water and potato tuber tissue. Tailocin production was induced in rejuvenated cells by the addition of 0.1 µg mL^-1^ mitomycin C (Abcam, Poland) or 0.003 % (0.88 mM) hydrogen peroxide (Laboratorium Galenowe Olsztyn Sp. z o. o., Poland) and cultures with no inductive factor were used as controls. Twenty-four hours post-induction, tailocins were PEG-purified from 2 mL volumes of filtered (0.2 µm, PES membrane) culture supernatants and resuspended in 1/10 of the initial volume in PBS (Borowicz et al., 2023). The concentration of tailocins was estimated by a semiquantitative spot test (Yao et al., 2017;Borowicz et al., 2023) using tailocin-susceptible strains (Supplementary Table S2). The reciprocal of the highest dilution causing a visible plaque was defined as the relative activity in arbitrary units (=1 AU), and the AU per mL of culture was calculated. The experiment was conducted twice, with tree replicates per combination.

### Competition assay *in vitro*

Competition assays were performed using a spontaneous rifampicin-resistant mutant of *D. solani* A623-S20-A17 selected on TSA with 50 µg mL^-1^ of rifampicin (Sigma-Aldrich)(Glandorf et al., 1992;Berg et al., 2007) and designated *Ds* RIF. This strain was tested in an *in vitro* competition assay against three sets of competitor strains. Each set comprised two *Dickeya* spp. strains which either: 1) were killed by the tailocins of *D. solani* A623-S20-A17 (*D. dianthicola* A260-S21-A16 and *D. zeae* A661-S21-A17), 2) produced tailocins killing *D. solani* A623-S20-A17 (*D. zeae* A586-S18-A17 and *D. oryzae* A3-S1-M15) or 3) were killed by the tailocins of *D. solani* A623-S20-A17 but at the same time produced tailocins targeting this strain (mutual killing)(*D. dadantii* 3937 and *D. chrysanthemi* A604-S21-A17). All strains were grown overnight in 0.1 TSB medium. Next, the cells were harvested by centrifugation (3500 RCR, 5 min.), and the pellet was resuspended in a fresh medium to obtain a turbidity of 0.5 McF. This equaled, on average, 1.4 × 10^8^ CFU ml^-1^ (±4.5 × 10^7^), verified by dilution plating. 150 µl of 0.1 TSB in a single well of a 96-well plate was inoculated (1:1, 25:25 µl) with the suspension of *Ds* RIF and one of the competing strains to ensure the strains had an even start and avoid tailocins carryover from the initial overnight cultures. Each co-culture was tested under two conditions: 0.1 TSB alone or 0.1 TSB with 0.003 % (0.88 mM) H_2_O_2_ (tailocin-inductive stress). The cultures were incubated for 22 h at 28°C, with orbital shaking, in the EPOCH2 reader (BioTek). Culture from each well was dilution-plated on two media types: TSB agar (for counting the total number of CFUs) and TSB agar with 50 µg mL^-1^ of rifampicin (for counting the fraction of *Ds* RIF). *Ds* RIF was tested against the wild-type (*wt*) variant of the strain as a control of the mutant’s fitness under the applied experimental conditions. Moreover, the ability of *Ds* RIF to produce tailocins was verified in a spot assay, as described above. The competition experiment was performed twice, with two separate co-cultures (replicates) per experiment.

## Results

### Most *Pectobacterium* spp. and *Dickeya* spp. strains produce phage tail-like particles

TEM imaging of preparations obtained from mitomycin C-induced cultures revealed that 22 out of 25 studied SRP isolates (88%) produce phage tail-like particles (tailocins) (Fig.1). The three exceptions were *Pectobacterium brasiliense* A143-S20-M16, *Pectobacterium atrosepticum* A597-S4-A17, and *Dickeya dianthicola* A260-S21-A16. All tailocins had similar morphologies in TEM and AFM images, regardless of whether they were derived from *Dickeya/Musicola* or *Pectobacterium* spp. (Fig. 1 and Fig. 2). The average length of tailocins produced by *Pectobacterium* spp. was 163±15 nm. Tailocins produced by *Dickeya/Musicola* spp. were significantly shorter, measuring 153±20 nm (Mann-Whitney test, p<0.05). Similarly, tailocins produced by *Pectobacterium* spp. were, on average, 2 nm wider than those of *Dickeya/Musicola* species, with the dimensions of 26±2 nm and 24±2 nm, respectively, with the difference being statistically significant (Mann-Whitney test, p<0.05). Three-dimensional images generated based on AFM scans for seven representative tailocin particles are shown in Fig. 2.

**Fig. 1.**
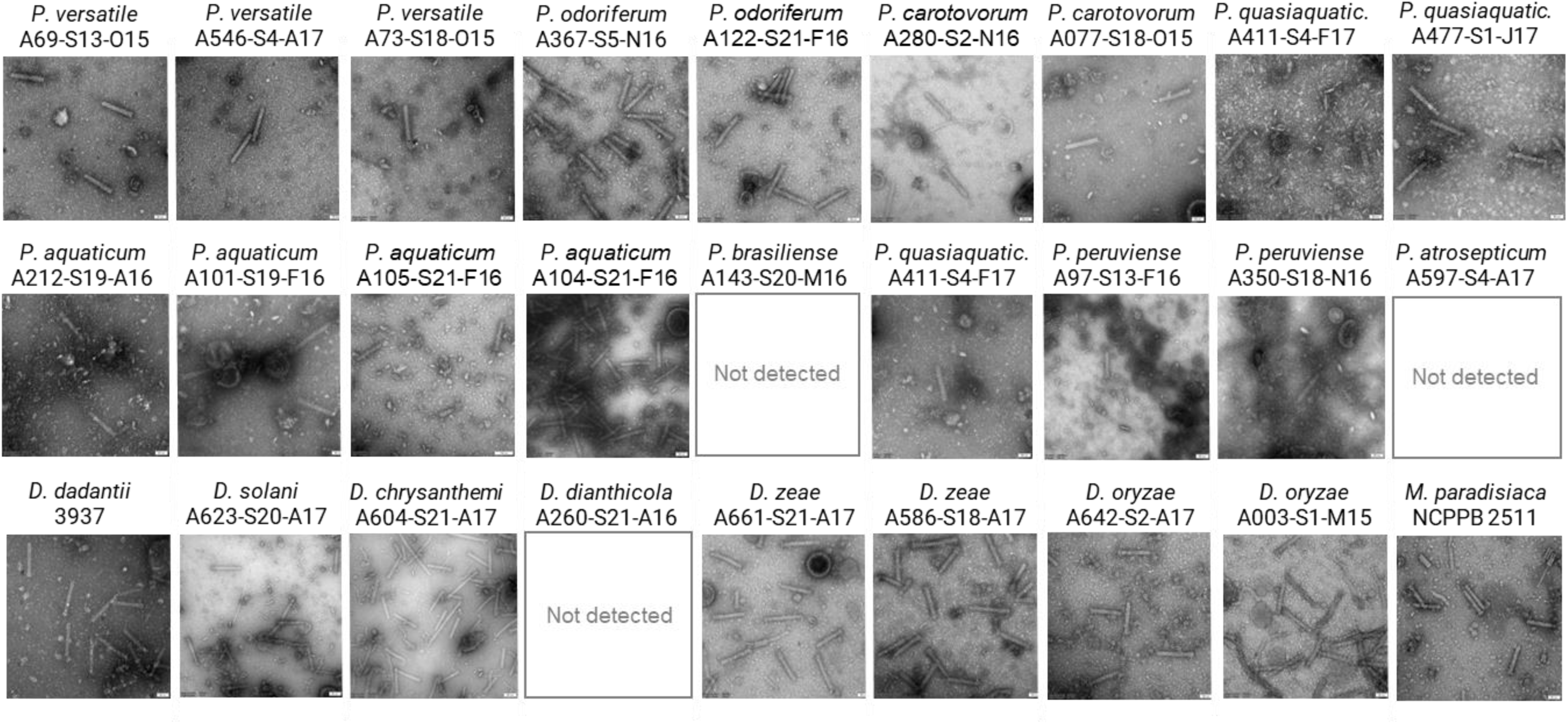
TEM images picturing tailocins isolated from the 27 analyzed SRP strains.

**Fig. 2.**
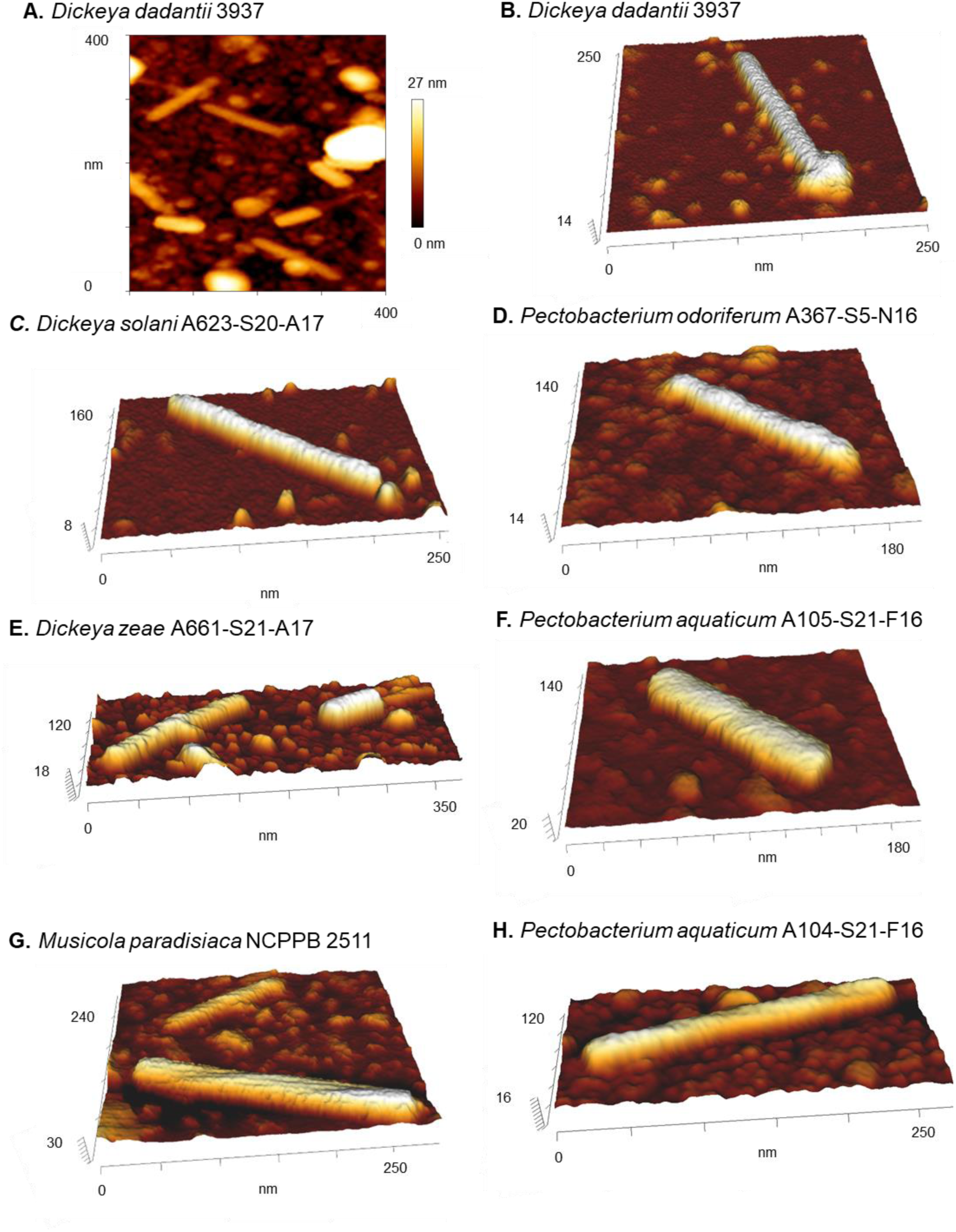
Images of the representative tailocins generated based on AFM scans. AFM imaging was performed using the JPK NanoWizard® 4 (NanoScience) in QI™ in the quantitative imaging (QI)™ force spectroscopy mode. Images were processed using JPK Data Processing software. Panel A shows the AFM scan of Dickeyocin P2D1 produced by *D. dadant*ii 3937, panels B-H. show the highest quality 3D visualizations of representative tailocin particles, generated based on AFM scans, for the following strains: *D. dadantii* 3937*, D. solani* A623-S20-A17, *D. zeae* A661-S21-A17. *M. paradisiaca* NCPPB 2511, *P. odoriferum* A367-S5-N16, *P. aquaticum* A105-S21-F16, *P. aquaticum* A104-S21-F16.

### The susceptible targets of SRP phage tail-like particles vary depending on the producing strain, with several exhibiting inter-genus toxicity

We evaluated the breadth of strains killed by phage tail-like particles produced by the SRP strains. In total, 702 unique tailocin-mediated interactions (producer-target combinations) were examined for the 351 unique pairs (individual producer vs. individual target) of SRPs tested (Fig. 3). The incidence of a strain being susceptible to a particular tailocin was 16.8% (Table 1). This killing incidence was higher among *Dickeya* strains than among *Pectobacterium* strains (57.1% and 21.6%, respectively) (Table 1). Strains *P. versatile* A73-S18-015 and *P. polaris* A641-S16-A17 were particularly susceptible to tailocins, being killed by those from 8 different strains. In contrast, *P. carotovorum* A280-S2-N16 was resistant to tailocins produced by all strains tested. Twenty-one of the 25 strains produced tailocins that killed at least one strain from the 27 SRPs. Of the 4 strains not exhibiting any killing activity, three (*P. brasiliense* A143-S20-M16, *P. atrosepticum* A597-S4-A17, and *D. dianthicola* A260-S21-A16) lacked visible tailocins in TEM analyses (Fig. 1), while *P. peruviense* A350-S18-N16 produces tailocins (as evidenced by TEM microscopy), these phage tail-like particles did not target any of the 26 SRPs tested (Fig. 3). In contrast, *D. oryzae* A3-S1-M15 and *P. odoriferum* A122-S21-F16 produced tailocins with the broadest target range, each killing 9 out of the 26 strains (33.3%).

**Fig. 3.**
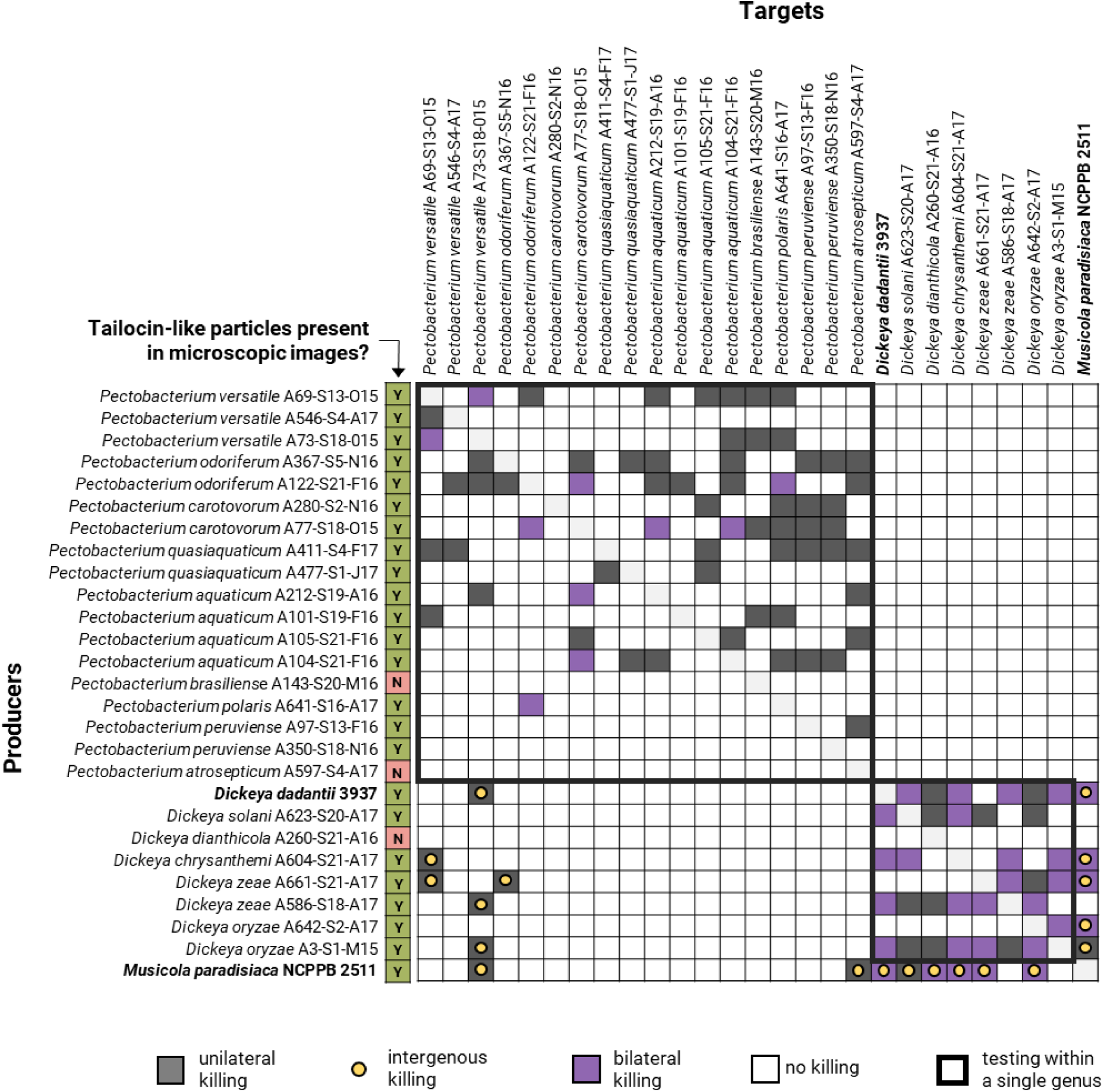
Matrix of tailocin-mediated interactions within the group of tested SRP strains. The tested pool comprised 27 strains: 25 isolates from the Durance River water and two type strains: *D. dadantii* 3937 and *M. paradisiaca* NCPPB 2511 (in bold). Grey-shaded cells indicate the susceptibility of a target strain to the tailocins of a given producer strain (unilateral killing). Unilateral killing where the producer and the target belong to different genera, is additionally marked by a yellow dot. Lillac cells also indicate the susceptibility of the target strain to the investigated tailocin, but in particular situations where the target strain was observed to produce ‘retributive’ tailocins against the producer (mutual killing). A dedicated column indicates whether tailocins were (Y) or were not (N) detected by microscopic imaging in culture supernatants of the tested strains following treatment with mitomycin C.

**Table 1.** Tailocin-mediated killing events for the tested pool of 27 SRP strains.

| Group | Total interactions<sup>A</sup> | Killing events |  | Mutual killing |  |  |
| --- | --- | --- | --- | --- | --- | --- |
|  |  | No. | % total | No. | % total | % killing events |
| All SRPs | 702 | 118 | 16.8% | 40 | 5.7% | 33.9% |
| *Pectobacterium* spp.<sup>B</sup> | 306 | 66 | 21.6% | 10 | 3.3% | 15.2% |
| *Dickeya* spp.<sup>C</sup> | 56 | 32 | 57.1% | 21 | 37.5% | 65.6% |
| Mixed genera<sup>D</sup> | 340 | 19 | 5.6% | 9 | 2.6% | 47.37% |
| *Psp* vs *Dsp* | 162 | 0 | 0% | 0 | 0% | 0% |
| *Dsp* vs *Psp* | 162 | 6 | 3.7% | 0 | 0% | 0% |
| *Dsp* vs *M. par.* | 8 | 5 | 62.5% | 4 | 50% | 80% |
| *M. par* vs *Dsp* | 8 | 6 | 75% | 5 | 62.5% | 83.3% |
| *M. par* vs *Psp* | 18 | 2 | 11.1% | 0 | 0% | 0% |
| *Psp* vs *M. par* | 18 | 0 | 0% | 0 | 0% | 0% |

<sup>A</sup> total interactions - the number of combinations where each strain from a subset was tested both for the potential to produce tailocins and for tailocin susceptibility (potential target)
<sup>B</sup> both strains in the tested interaction assigned to *Pectobacterium* spp. (total of 18 strains tested)
<sup>C</sup> both strains in the tested interaction assigned to *Dickeya* spp. (total of 8 strains tested)
<sup>D</sup> each strain of the pair representing a different genus; *Psp* – *Pectobacterium* spp.; *Dsp* – *Dickeya* spp.; *M.* *par* – *Musicola paradisiaca* NCPPB 2511. The first strain in each combination is the tailocin producer.

We also investigated the ability of tailocins to kill stains in genera different from the producer. *Pectobacterium* strains produced tailocins targeting only other members of the genus. In contrast, 5 out of 8 tested *Dickeya* spp. (62.5%) could kill at least one representative of *Pectobacterium* sp., either a *P. versatile* strain or the closely related *P. odoriferum. M. paradisiaca* NCPPB 2511, the single representative of *Musicola* spp., strongly resembled *Dickeya* spp. both in tailocin susceptibility and the specificity of its tailocins (Fig. 3, Table 1).

### Several SRP strains produced tailocins that targeted other tailocin-producing strains, resulting in a phenomenon known as ‘mutual killing’

Analysis of tailocin-mediated interactions among the collection of SRPs revealed that 33.9% of the killing events were retaliated (bilateral) – in which strains could kill each other (Table 1). For 20 of the 351 pairs of SRPs tested (5.7%), a given strain pair was mutually inhibitory - a phenomenon we designate ‘mutual killing’. Mutual killing was more common within *Dickeya* spp. (65.6% of killing events were mutual) than within *Pectobacterium* spp. (15.2%), and we observed no mutual killing between members of *Dickeya* and *Pectobacterium* (Table 1). The mutual killing was very common (80% of killing events) in mixed pairs of *Dickeya* strains and *M. paradisiaca* NCPPB 2511, reflecting the clustering between strain NCPPB 2511 and *Dickeya* spp.

### Tailocins of *Dickeya* spp. and *Pectobacterium* spp. are phylogenetically distinct

Based on the tailocin-associated protein identified by mass spectrometry, we identified gene clusters responsible for tailocin production in the genomes of the SRP strains. Among the 25 strains examined, tailocin-encoding clusters were absent only in 2 strains - *Pectobacterium brasiliense* A143-S20-M16 and *Pectobacterium atrosepticum* A597-S4-A17. These two strains, together with *Dickeya dianthicola* A260-S21-A16, also lacked tailocin production by imaging.

We also performed synteny analyses of the tailocin clusters within the *Pectobacterium* and *Dickeya* (including *Musicola*) genera (Fig. 4). The cluster in *Pectobacterium* sp. exhibited homology with that encoding carotovoricin Er (Nguyen et al., 2002) while those in *Dickeya* species and *Musicola* sp. were genetically similar to those encoding dickeyocin P2D1 (Borowicz et al., 2023). Furthermore, no SRP strain possessed both carotovorocin Er and P2D1 encoding clusters. *Pectobacterium* spp. strains exclusively contained the carotovorocin Er-like cluster, while *Dickeya* spp. strains solely harbored cluster encoding P2D1-like tailocin. Thus, no strain that contains both types of tailocins has been identified.

**Fig. 4.**
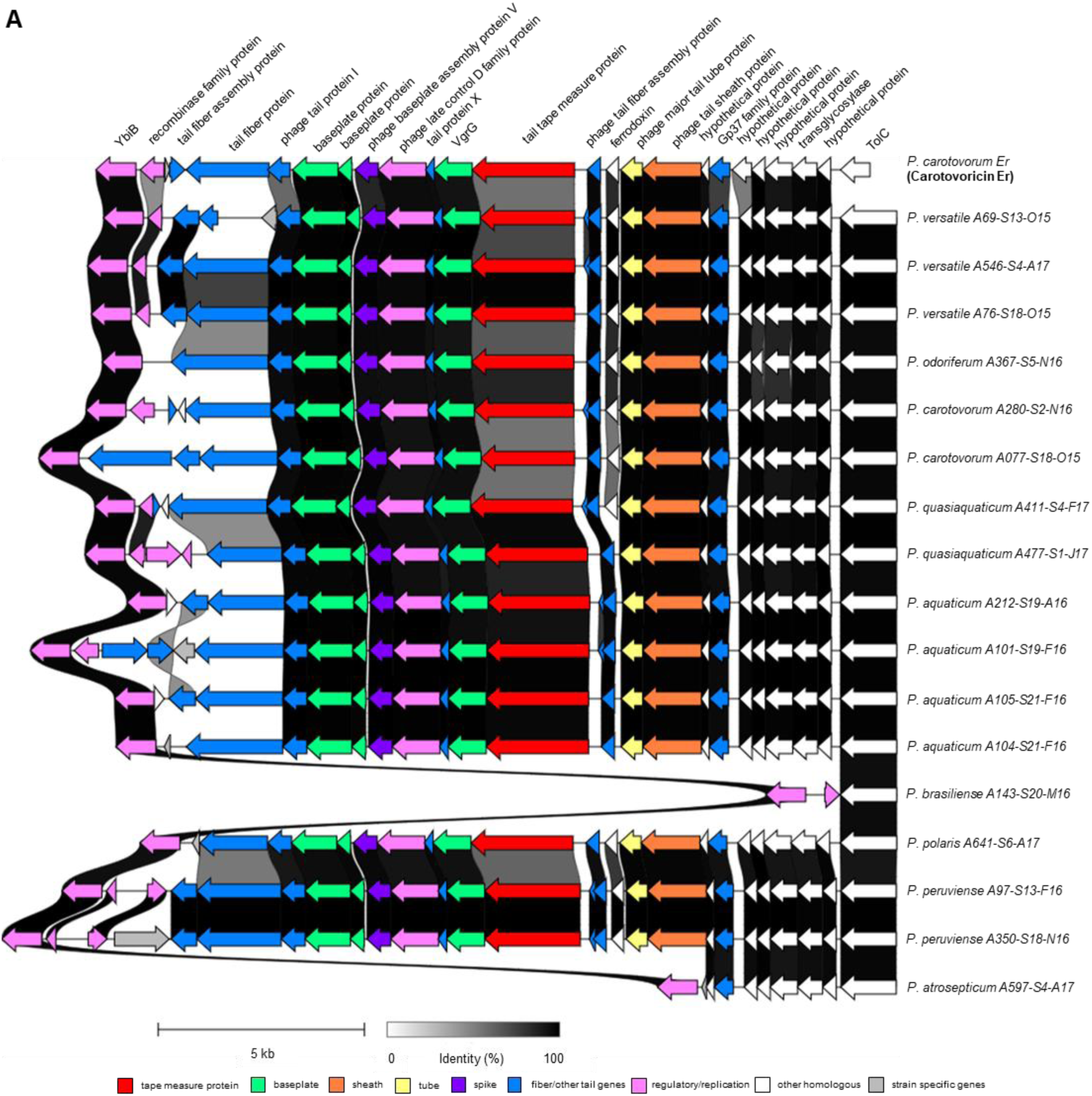
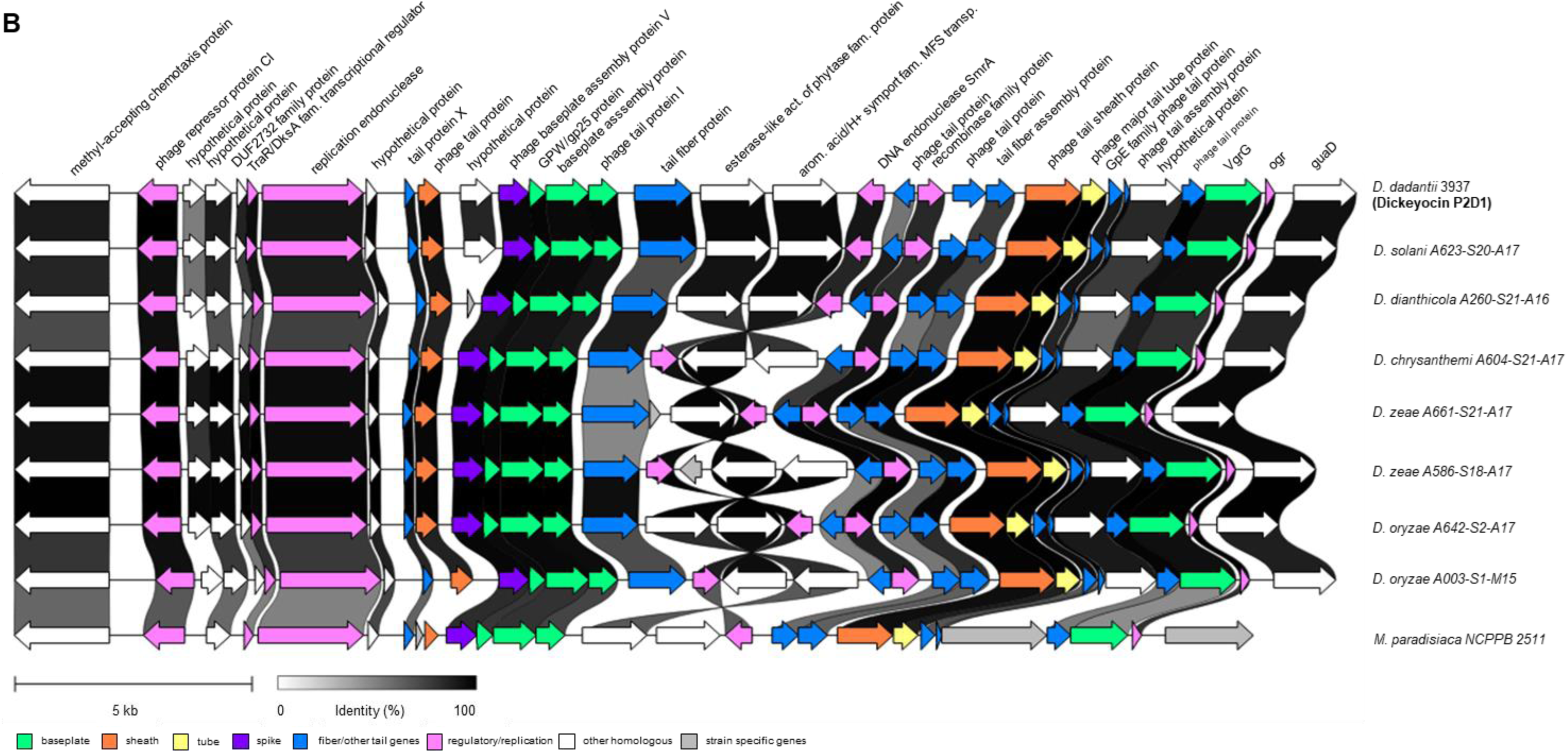
Comparison of tailocin clusters in the analyzed group of SRPs. The upper panel (A) depicts regions homologous to the carotovoricin-encoding cluster typical for *Pectobacterium* spp. (reference: carotovoricin Er accession: AB045036.1, *P. versatile* A69-S13-O15 RMBQ01000001.1, positions in the genome: 94551-113781). The lower panel (B) shows regions homologous to dickeyocin P2D1, typical for *Dickeya* spp. and also present in *Musicola paradisiaca* NCPPB 2511 (reference: *D. dadantii* 3937 NC_014500.1, positions in 0 the genome 2730482-2757061).

The overall genomic organization of the tailocin clusters was conserved within each genus (Fig. 4). In *Pectobacterium* species (Fig. 4A), the most prominent difference was the presence of a highly variable gene annotated as encoding a tail fiber protein/pyocin knob domain-containing protein, that appeared to contain segments shared among different strains (data not shown). Additional, albeit less pronounced, variability was observed in the sequence of the gene encoding a tail tape measure protein. In *P. brasiliense* A143-S20-M16, this cluster was flanked by genes conserved across other strains; however, instead of a complete tailocin cluster, it contained only the single gene annotated as a mobile element protein gene. In *P. atrosepticum* A597-S4-A17, this region included not only conserved flanking genes but also seven genes present in other clusters and one strain-specific gene. In *Dickeya* species (and *Musicola* sp.) (Fig. 4B), the main differences between strains included (1) the presence of strain-specific genes likely unrelated to tailocin production, (2) the orientation of genes not directly involved in tailocin structural components (such as aromatic acid/H+ transporters and phytase family proteins), (3) sequence variations in intergenic regions (data not shown), and (4) sequence differences in tail fiber genes. Notably, *D. dianthicola* A260-S21-A16, which did not produce tailocins, did not exhibit substantial differences in the genomic region encoding tailocins in other strains. Specifically, the tailocin cluster was present in the genome and showed high homology to that of other *Dickeya* spp. tailocin clusters.

### Carotovoricin and dickeyocin clusters are common in the genomes of *Pectobacterium* spp. and *Dickeya* spp

Given that not all SRPs examined produced tailocins nor harbored the tailocin cluster, we determined the prevalence of tailocin clusters across a broader range of SRP strains to assess the frequency of this trait better. We examined the complete genomes of 116 *Pectobacterium* spp. for the presence of the carotovoricin cluster and 74 genomes of *Dickeya* and 1 *Musicola* spp. genome for the presence of genes encoding dickeyocin. Within the *Pectobacterium* clade, 96 genomes (83%) harbored homologous tailocin regions, having >70% coverage and identity. Twelve *Pectobacterium* genomes had partial alignment (10% of strains), and 9 (8%) showed no evidence of the tailocin cluster (Dataset S1). Within the many *Dickeya* spp. and the single *Musicola* spp. genomes examined, 52 (69%) had a cluster that aligned well with the dickeyocin cluster, while 16 (21%) genomes had coverage below 70%, and 7 (9%) genomes had no apparent tailocin cluster (Dataset S2). None of the 6 complete genomes of the species *P. atrosepticum* present in the NCBI database had a tailocin cluster.

### Tailocin-killing specificity is not aligned with phylogenetic distance in SRP strains

The topology of the dendrograms of the phylogenetic tree of SRP based on MultiLocus Sequence Analysis (MSLA) was not well aligned with that based on the patterns of tailocin killing position (Supplementary Fig. S1). This suggests that tailocin activity may be influenced by factors other than taxonomic relationships.

### Constitutive and induced production of tailocins in *Dickeya* and *Pectobacterium* spp

Given that our previous study revealed the surprising apparent constitutive basal level of tailocin production in *D. dadantii* 3937 (Borowicz et al., 2023), we explored whether this was also the case in other strains. Tailocin production was thus assessed in 6 strains (3 *Pectobacterium* and 3 *Dickeya* spp.) (*P. versatile* A69-S13-O15, *P. quasiaquaticum* A411-S4-F17, *P. aquaticum* A212-S19-A16, *D. dadantii* 3937, *D. chrysanthemi* A604-S21-A17, *D. zeae* A586-S18-A17).

Furthermore, we wished to establish whether reactive oxygen species (ROS), such as H_2_O_2,_ often associated with plant infection (Torres et al., 2006;Reverchon et al., 2016), could induce the production of tailocins in SRPs. Basal levels of tailocin production were observed for all tested strains even in the absence of ROS. However, the basal level of tailocins produced varied substantially between strains, with the highest basal production observed for the model strain *D. dadantii* 3937 (7 × 10^4^ AU mL^-1^) and the lowest for *P. aquaticum* A411-S4-F17 (5 × 10^2^ AU mL^-1^) (Fig. 5). The average abundance of tailocin was slightly (43%) higher for *Dickeya* than for *Pectobacterium* spp. (Mann-Whitney test, p<0.05) (2.8 × 10^4^ and 1.6 × 10^4^ AU mL^-1^ of culture, respectively).

**Fig. 5.**
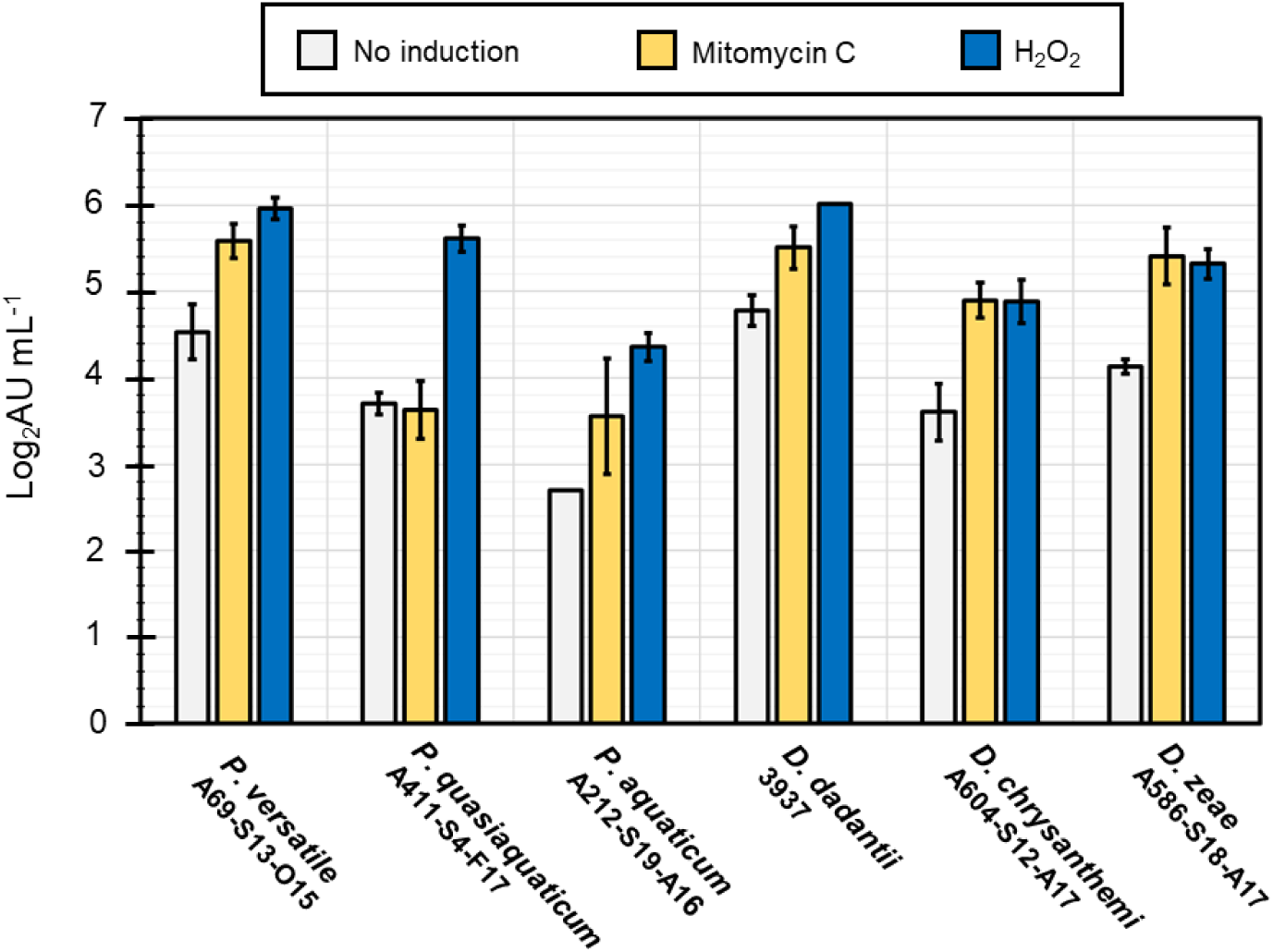
Basal and induced tailocin production levels in six *Pectobacterium* spp. And *Dickeya* spp strains. The graph shows the logarithm of the arbitrary tailocin units per mL of culture (Log AU mL^-1^) for three strains of *Pectobacterium* spp. (*P. versatile* A69-S13-O15, *P. quasiaquaticum* A411-S4-F17, *P. aquaticum* A212-S19-A16) and three strains of *Dickeya* spp. (*D. dadantii* 3937, *D. chrysanthemi* A604-S21-A17, *D. zeae* A586-S18-A17). The basal production levels (no induction) are compared to those with mitomycin C or hydrogen peroxide after induction. Averages from 2 experiments, each consisting of 3 separate inductions, are shown, and error bars represent the standard deviation.

Mitomycin C (0.1 µg mL^-1^, control) caused a significant induction (approx. 5 to 20-fold) of tailocin production in all strains except *P. quasiaquaticum* A411-S4-F17 (Fig. 5). The highest induction was observed for *D. zeae* A586-S18-A17 (19-fold) and *D. chrysanthemi* A604-S21-A17 (20-fold). Meanwhile, the induction of tailocins by mitomycin C in *D. dadantii* 3937 was relatively low (5-fold) due to its high production in the absence of mitomycin.

Hydrogen peroxide also induced tailocin production in all strains (approx. 15 to 81-fold) (Fig. 5). For most strains, hydrogen peroxide (0.88 mM, 0.003%) conferred a higher yield of tailocins than that induced by mitomycin C (0.1 µg mL^-1^). Notably, while no induction of tailocins by mitomycin in *P. quasiaquaticum* A411-S4-F17 was observed, the addition of hydrogen peroxide resulted in the highest increase compared to basal production levels of all strains (81-fold) (Fig. 5).

### Detectable levels of phage tail-like particles are produced in rotting potato tuber tissue but not in river water

Tailocin production was explored in SRP in a variety of natural settings in addition to cultural media. Three *Pectobacterium* spp: *P. versatile* A69-S13-O15, *P. quasiaquaticum* A411-S4-F17, *P. aquaticum* A212-S19-A16, and three *Dickeya* spp.: *D. dadantii* 3937, *D. chrysanthemi* A604-S21-A17, *D. zeae* (very recently reclassified as *D. parazeae* (Hugouvieux-Cotte-Pattat and Van Gijsegem, 2021)) (A586-S18-A17) were inoculated into river water (the source of the strains) as well as into potato tubers. Tailocin production was not detected in any strain in river water (Table 2) but was abundant in macerated potato tuber tissue. Tailocins were found not only in the symptomatic tissues caused by those strains that were pathogenic to potato (*P. versatile* A69-S13-O15, *D. dadantii* 3937, *D. chrysanthemi* A604-S21-A17, *D. zeae* A586-S18-A17) but also in healthy tuber tissue surrounding the site where non-pathogenic strains (*P. aquaticum* A212-S19-A16 and *P. quasiaquaticum* A411-S4-F17) were inoculated (Table 2) (Supplementary Fig. S2).

**Table 2.**
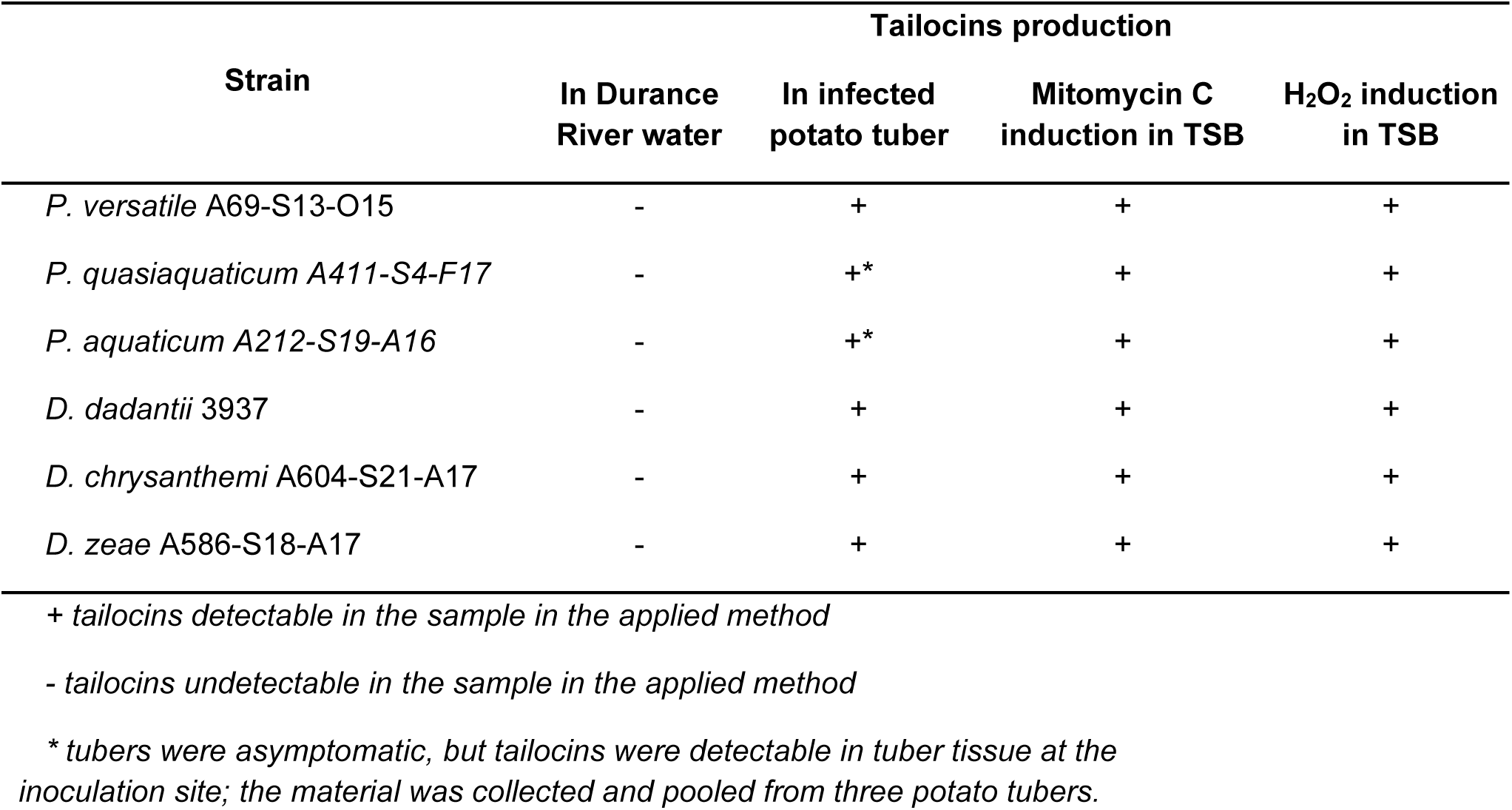
Production of tailocins in river water and potato tubers. Qualitative tailocin production in different conditions for various bacterial strains. The strains tested include *P. versatile* A69-S13-O15, *P. quasiaquaticum* A411-S4-F17, *P. aquaticum* A212-S19-A16, *D. dadantii* 3937, *D. chrysanthemi* A604-S21-A17, and *D. zeae* A586-S18-A17. Tailocin production was assessed in infected potato tubers, in Durance River water, and as a positive control following Mitomycin C and H_2_O_2_ induction in TSB (Tryptic Soy Broth). The symbols represent the following: “+” indicates detectable tailocins in the sample with the applied method; “-” indicates undetectable tailocins in the sample with the applied method; “*” indicates that tailocins were detectable if tuber tissue in direct contact with inoculated bacteria was pooled from three potato tubers.

| Strain | Tailocins production |  |  |  |
| --- | --- | --- | --- | --- |
|  | In Durance River water | In infected potato tuber | Mitomycin C induction in TSB | H<sub>2</sub>O<sub>2</sub> induction in TSB |
| *P. versatile* A69-S13-O15 | - | + | + | + |
| *P. quasiquaticum* A411-S4-F17 | - | + | + | + |
| *P. aquaticum* A212-S19-A16 | - | + | + | + |
| *D. dadantii* 3937 | - | + | + | + |
| *D. chrysanthemi* A604-S21-A17 | - | + | + | + |
| *D. zeae* A586-S18-A17 | - | + | + | + |

+ tailocins detectable in the sample in the applied method
- tailocins undetectable in the sample in the applied method
\* tubers were asymptomatic, but tailocins were detectable in tuber tissue at the inoculation site; the material was collected and pooled from three potato tubers.

### Tailocin production is not always sufficient to ensure competitive success against susceptible neighbors

We explored the extent to which tailocin production by *D. solani* A623-S20-A17 (*Ds*) mediated the outcome of two-strain co-cultures with strains susceptible to the strain’s tailocins or/and was targeted by tailocins of the competitor (Fig. S3). To monitor changes in the proportion of *Ds* in mixed cultures, we selected a spontaneous rifampicin-resistant mutant of *Ds* (*Ds* RIF), which could be selectively recovered on rifampicin-containing media. Ds RIF retained tailocin production and did not differ from the parental strain in growth rate in 0.1 TSB or ability to compete with the wt *in vitro*. These results indicate that the rifampicin*-*resistant *Dickeya* spp. strains are not compromised in competitiveness under the applied experimental conditions (Fig. 6). The competitive interactions were explored with three types of competitors: 1) those susceptible to the tailocins of *Ds* RIF, 2) those producing a tailocin inhibitory to *Ds* RiF, or 3) those for which mutual killing with Ds RIF occurs in vitro. The success of a strain was defined as the extent of its end-point dominance in the co-culture (>90% of total CFU) following its initial inoculation in equal proportions (Fig. S3). Two competitors that could kill *Ds* RIF with their tailocins prevailed over *Ds* RIF, suggesting that tailocins contributed to this success. However, Ds RIF outcompeted only a single strain that was susceptible to its tailocin and only in the presence of H_2_O_2_. This suggests that tailocins alone do not guarantee the success of a producer against a susceptible rival.

**Fig. 6.**
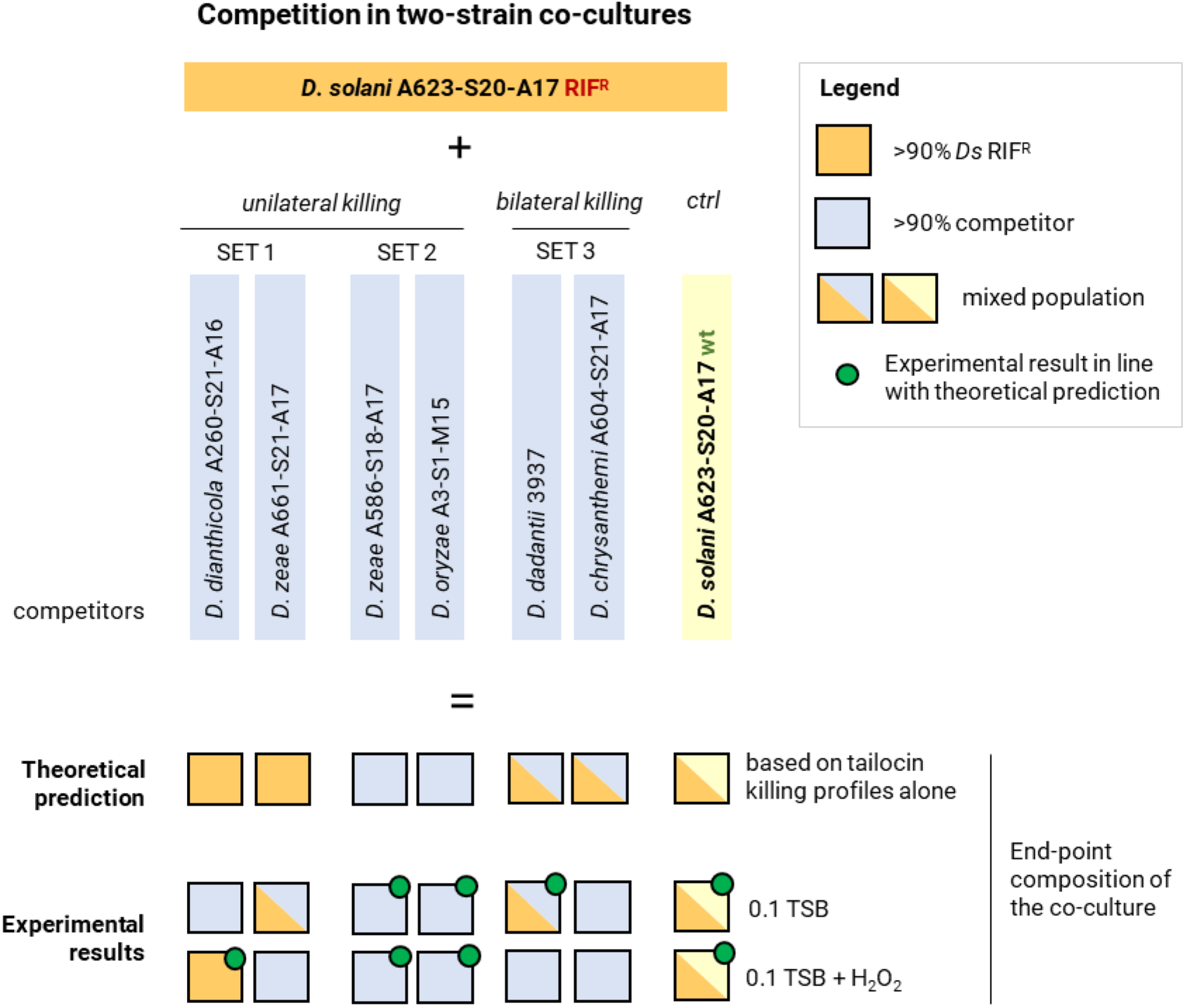
Competition between *D. solani* A623-S20-A17 RIF^R^ (*Ds* RIF) and other SRP strains in two-strain co-cultures. The success was defined as the end-point dominance of a given strain in the co-culture (>90% of total CFU), with both strains ensured with an even start (1:1 mix; ca. 10^7^ CFU/ml/strain). The theoretical winners, predicted based on tailocin-mediated interactions, were compared with those that were experimentally established. Co-cultures were conducted for 22 h in 0.1 TSB alone or supplemented with 0.88 mM H_2_O_2_ (tailocin-inductive conditions). SET 1: strains sensitive to tailocins of *Ds* RIF. SET 2: strains producing tailocins against *Ds* RIF. SET 3: strains sensitive to tailocins of *Ds* RIF and producing tailocins targeting this strain (mutual killing). *ctrl* – control experiment; competition between *Ds* RIF and the *wt* variant of this strain.

## Discussion

Our study revealed that most SRP strains produce phage tail-like particles. While *P. brasiliense* A143-S21-A16 and *P. atrosepticum* A597-S4-A17 lacked tailocin genomic clusters, *D. dianthicola* A260-S21-A16 had a complete cluster but did not produce tailocins. This could be due to reliance on other competitive strategies (Hibbing et al., 2010) or regulation by specific environmental factors not present in the experiment (Bellieny-Rabelo et al., 2019). Furthermore, we assessed the prevalence of tailocin clusters in 190 high-quality, complete genomes of SRP bacteria and found that 83% of *Pectobacterium* spp. and 69% of *Dickeya* spp. genomes contain clusters encoding tailocins. This indicates that tailocins are a very common feature among SRP and may suggest that their production contributes to the ecological success of these bacteria (Ma et al., 2007). The prevalence of tailocins in SRP aligns with the previous observations documenting their presence in various bacterial species, including *Pseudomonas* spp. (Stice et al., 2023), *Pantoea* spp. (Nagakubo et al., 2021) and *Streptomyces* spp. (Nagakubo et al., 2021).

Most tailocins can kill only closely related strains. This finding has, however, usually been based on studies of a small set of strains either at the species or genus level (Granato et al., 2019). Some observations suggest that some phage tail-like particles can kill different species than the producer (Yao et al., 2017;Principe et al., 2018). Our work interrogated a large group of strains belonging to three genera for their mutual tailocin-based interactions. While some SRP strains produce tailocins with a broad range of activity, killing more than 30% of other strains of different species. Caution, therefore, should be made to accept the current dogma of a narrow killing range of tailocins.

Interestingly, while some *Dickeya* spp. produced tailocins able to kill *Pectobacterium* spp. cells, the opposite was not observed in our study. Due to their evolutionary association with bacteriophage tails (Ghequire and De Mot, 2015), it has been suggested that tailocins might retain some degree of flexibility in their host range, such as known phage-host interactions (Backman et al., 2024a). Inter-genus activity may remain an important trait in competition as mixed bacterial communities involving both genera are frequently observed in the field, and the ability to target a broader range of competitors could confer a selective advantage (Perombelon, 1988;Charkowski, 2009;2018;Barny et al., 2024).

Likewise, the observation of mutual killing, where two strains reciprocally target each other with tailocins, has not been extensively documented. The higher incidence of mutual killing within *Dickeya* spp. compared to *Pectobacterium* spp. suggests some (yet unknown) ecological factors may select for this behavior more often in *Dickeya* spp. than in *Pectobacterium* spp. This is somehow surprising, given that multiple species of SRP bacteria are frequently found together in the same environments (Toth et al., 2021a;van der Wolf et al., 2021). In such settings, these various species would be expected to experience similar ecological pressure and benefit from reciprocal tailocin production (Shyntum et al., 2019).

Phage tail-like particles isolated from both SRP genera exhibited similar morphological characteristics despite the apparent phylogenetic differences in the origins of tailocin clusters in *Pectobacterium* spp. and *Dickeya* spp. suggests that these phage tail-like particles have been acquired independently in those genera. All of the clusters present in *Dickeya* spp. exhibit homology to that encoding P2D1 tailocin of *D. dadantii* 3937 (Borowicz et al., 2023), whereas all clusters present in *Pectobacterium* spp. express homology to the cluster encoding carotovoricin Er (Yamada et al., 2006). The fact that these phage tail-like particles were observed in all *Dickeya* spp. and most of the *Pectobacterium* spp. strains tested suggest that the tailocin clusters were acquired before species diversification in both genera and were subsequently maintained rather unaltered. As tailocins are large protein complexes and their synthesis should be a burden for a producer cell, their conservation suggests that tailocin production plays an important role in the ecology of these bacteria (Glasner et al., 2008;Hugouvieux-Cotte-Pattat, 2016).

The genes encoding tailocin fiber proteins exhibited the lowest homology in the tailocin clusters both in *Dickeya* spp. and *Pectobacterium* spp. This confirms that fiber proteins play a pivotal role in determining the specificity of tailocins and their killing range, as reported earlier (Dams et al., 2019). Fiber proteins are the most important determinants of target recognition in bacterial cells (Backman et al., 2024a). This specificity is crucial for the success of tailocins as killing agents, enabling them to discriminate between susceptible and resistant bacterial strains. In bacteriophages, receptor-binding proteins are known to undergo positive selection to adapt to variations in surface receptors of target bacteria, leading to high variability even among closely related strains (Bertozzi Silva et al., 2016). In SRP, the killing spectrum of tailocins, determined by the sequence of fiber proteins, appears to be more dependent on ecological factors and the pressure of host-pathogen interactions encountered by the individual strains than the evolutionary relationships between them.

Although mitomycin C is widely used to trigger tailocin induction under laboratory conditions, in natural and agricultural environments, the likely inducers of tailocins remain unknown. It seemed likely that other factors that cause DNA damage and thus activate the SOS pathway could also serve as inducers of tailocins. In the proof-of-concept experiment, we found that hydrogen peroxide was a potent inducer of SRP phage tail-like particles. Like mitomycin C, hydrogen peroxide is known to activate the ROS response (Storz et al., 1990;Erill et al., 2007). During infection, SRP bacteria often encounter oxidative stress in plant hosts during the immune response of the plant (Reverchon and Nasser, 2013;Jiang et al., 2016). Therefore, the induction of tailocins by reactive oxygen species may serve as a mechanism for bacterial survival in their natural settings, aiding in eliminating competing bacteria under stressful conditions. Indeed, the highest levels of competition might be expected during the infection process, where the change in the plant nitch during injection leads to very high bacterial numbers. The ability of plant pathogenic bacteria, including SRP, to constitutively produce tailocins, as well as under stressful conditions, such as exposure to ROS, may lead to important ecological consequences. The basal tailocin production by a fraction of the SRP populations may be seen as an altruistic behavior in which a small number of bacterial cells sacrifice themselves to confer an advantage for the entire population over other microorganisms residing in the same location (Griffin et al., 2004). Alternatively, basal tailocin production may be understood as a division of labor in which subpopulations of bacteria are ‘assigned’ to different tasks to grant success to the entire population in a given niche (Lee et al., 2010;West and Cooper, 2016;Zhang et al., 2016). The ability to express tailocins in response to such oxidative stress might give SRP a competitive edge, allowing them to eliminate competitors in the plant environment when they must compete with all other microorganisms (Backman et al., 2024b).

The variable level of response to mitomycin C and hydrogen peroxide we observed in SRP strains highlights the still unraveled complexity of tailocin regulation. This suggests that in SRP, depending on the strain and type of inducer, the production of tailocins is regulated differently in different strains. Similar observations were made for prophages, where various environmental factors induced phages with different efficiency (Casjens, 2003;Varani et al., 2013).

The SRP strains addressed in this study were isolated from a single habitat. Despite this, this collection comprised both aquatic and non-aquatic strains, most of which (except *P. aquaticum* and *P. quasiaquaticum* strains known to be avirulent on plants) infect plants in agricultural settings (Ben Moussa et al., 2022). We found that tailocins are produced in infected potato tubers but not in river water. This aligns with the other studies in which tailocins were found to be produced primarily in nutrient-rich, competitive environments (Carim et al., 2021). Decaying plant material would be expected to be a much more nutrient-rich environment than river water. The environmental conditions in potato tubers might provide several stimuli for tailocin production, including ROS, plant exudates, cell wall degradation products, and microbial competition. In contrast, water is neither the common niche nor the environment where competition is likely to be intense. Instead, water rather serves as a translocation medium from one host to another (Pédron and Van Gijsegem, 2019;van der Wolf et al., 2021). This observation may be particularly pertinent for SRP bacteria, which compete with each other and other microorganisms for resources present within the plant tissues (Reverchon and Nasser, 2013). The potato tuber tissue represents a primary ecological niche for SRP pathogens (Perombelon and Kelman, 1980;Pérombelon and Salmond, 1995). Consequently, the presence of tailocins in both rotting tissue and the healthy surrounding tissue suggests that tailocin production may be part of the virulence strategy to minimize the risk of competitive infections.

Surprisingly, SRP strains belonging to species that are not found on plants (Portier et al., 2020) and which do not provoke disease symptoms (*vide*: *P. quasiaquaticum* A411-S4-F17 and *P. aquaticum* A212-S19-A16) did produce tailocins in the plant environment but not in river water. The low nutrient availability, the cost of production of this complex machinery, and the few direct competitors of SRP bacteria in river water likely explain the absence of tailocin production in this environment (Lamichhane and Bartoli, 2015) (Bradford et al., 2013;Toth et al., 2021b). The fact that these tailocin clusters were maintained in these water-associated species and were induced by the same stress trigger as plant-associated species suggests that tailocin production is not solely tied to SRP virulence on plants but could also play a role in other niches where these bacteria encounter stress. For example, it has been suggested that these water-associated species could be associated with insect larvae in the water stream (Ben Moussa et al., 2022). Whether tailocin production is triggered in such an environment remains to be determined.

Multiple factors play roles in the competition between microbes, with success being the sum of factors/traits of varying importance depending on the environmental context (Hibbing et al., 2010). In our study of mixed populations of tailocin producers and targets, tailocins were not always sufficient to ensure the dominance of producers over susceptible strains. The widespread occurrence of tailocin clusters among SRP led us to conclude that tailocin-mediated success is often conditional. It is well known that SRP bacteria use diverse weaponry for intra-genus and intra-genus interactions; therefore, tailocin production may be just one possibility to compete with kin microbes.

Several factors, such as environmental conditions, strain-specific interactions, and the relative abundance of producers versus susceptible strains, can influence the outcome of competitive interactions. This study explored an ‘even start’ scenario, where tailocin-producing and susceptible competitors were initially mixed in equal cell numbers. In bacterial populations, even when one strain has an antagonistic advantage (like producing phage tail-like particles), an equal start condition can neutralize this advantage because the susceptible strain is not immediately overwhelmed by that trait (Ross-Gillespie and Kummerli, 2014). Moreover, tailocin production, unlike the secretion of most bacterial antimicrobials, requires the lysis of the producer cells. Ergo, it can negatively influence the growth rate of the producer’s population. This burden can be assumed to be negligible when the producer-to-target ratio is high. Therefore, we speculate that tailocin production may be useful to SRP bacteria to protect already conquered niches from being invaded by competitors, a process named niche exclusion (Bauer et al., 2018).

## Author Contributions

MB: Conceptualization, Investigation, Methodology, Visualization, Writing – original draft, Writing – review & editing, DMK: Conceptualization, Investigation, Supervision, Visualization, Writing – original draft, Writing – review & editing, MS: Investigation, Methodology, Visualization, MN: Investigation, Methodology, Visualization, Writing – original draft, IM: Investigation, Methodology, Writing – original draft, PC: Methodology, Writing – original draft, JP: Conceptualization, Investigation, Methodology, Writing – review & editing, MAB: Conceptualization, Investigation, Methodology, Writing – review & editing, PYC: Investigation, Methodology, JD: Conceptualization, Investigation, Methodology, RC: Conceptualization, Data curation, Funding acquisition, Resources, Supervision, Writing – original draft, Writing – review & editing

## Data availability statement

All data can be obtained from the corresponding author and will be shared freely upon reasonable request.

## Acknowledgments

This research was financially supported by the National Science Center, Poland (Narodowe Centrum Nauki, Polska) via a research grant SONATA BIS 10 (2020/38/E/NZ9/00007) to Robert Czajkowski. The authors would like to express their gratitude to Prof. Steven E. Lindow (University of California-Berkeley, Berkeley, CA, United States) for his comments on the manuscript and his editorial work.

## Supplementary Tables

**Table S1.**
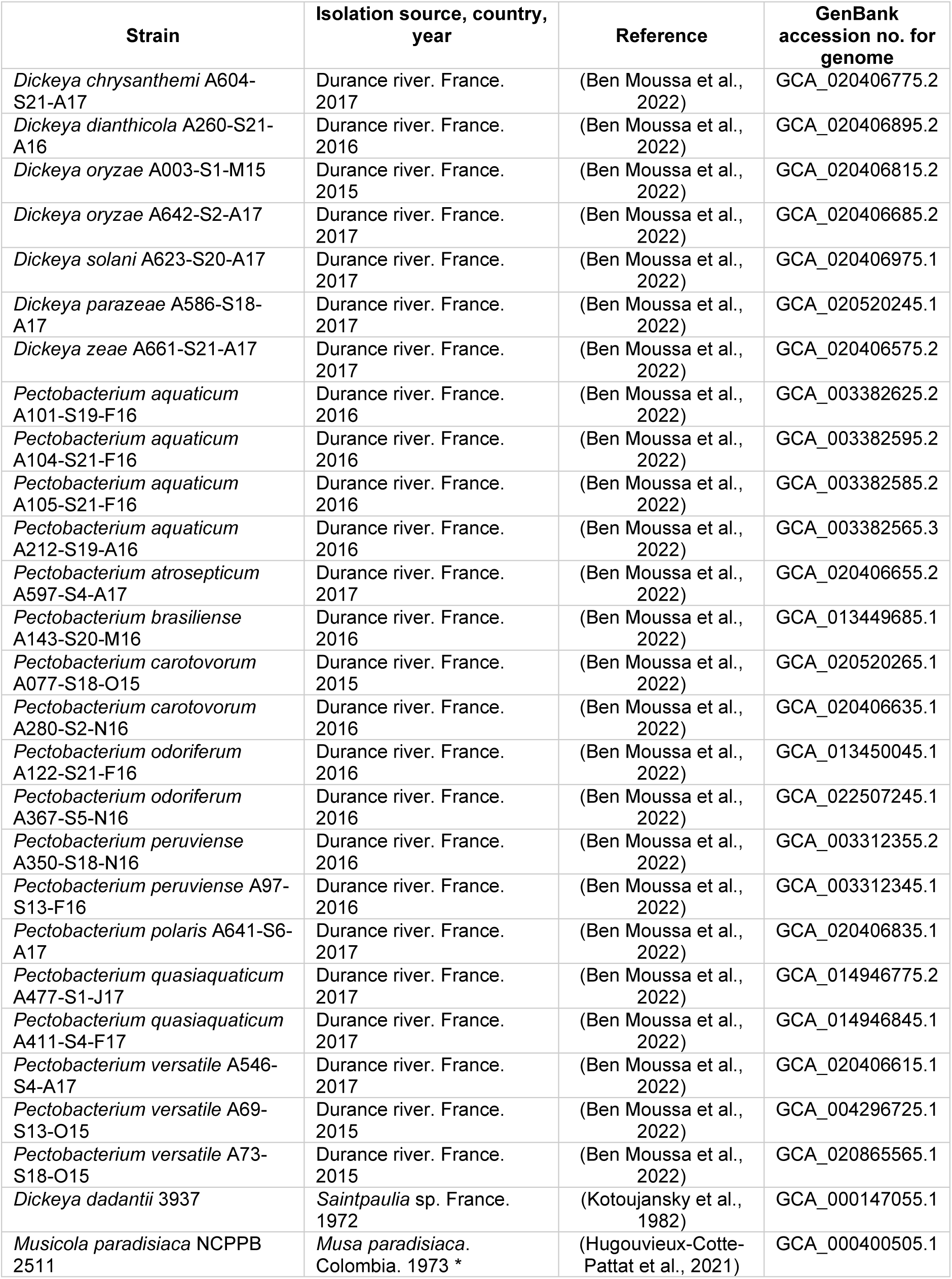
List of bacterial strains used in this study.

**Table S2.** List of sensitive strains used in a spot assay to detect tailocins from the given producer strains.

| **Tailocin producer** | **Sensitive strain** |
| --- | --- |
| *P. versatile* A69-S13-O15 | *P. aquaticum* A105-S21-F16 |
| *P. quasiquaticum* A411-S4-F17 | *P. atrosepticum* A597-S4-A17 |
| *P. aquaticum* A212-S19-A16 | *P. aquaticum* A105-S21-F16 |
| *D. dadantii* 3937 | *M. paradisiaca* NCPPB 2511 |
| *D. chrysanthemi* A604-S21-A17 | *D. zeae* A586-S18-A17 |
| *D. zeae* A586-S18-A17<br>(recently reclassified to *D. parazeae*) | *M. paradisiaca* NCPPB 2511 |

## Supplementary Figures

**Supplementary Fig. S1.**
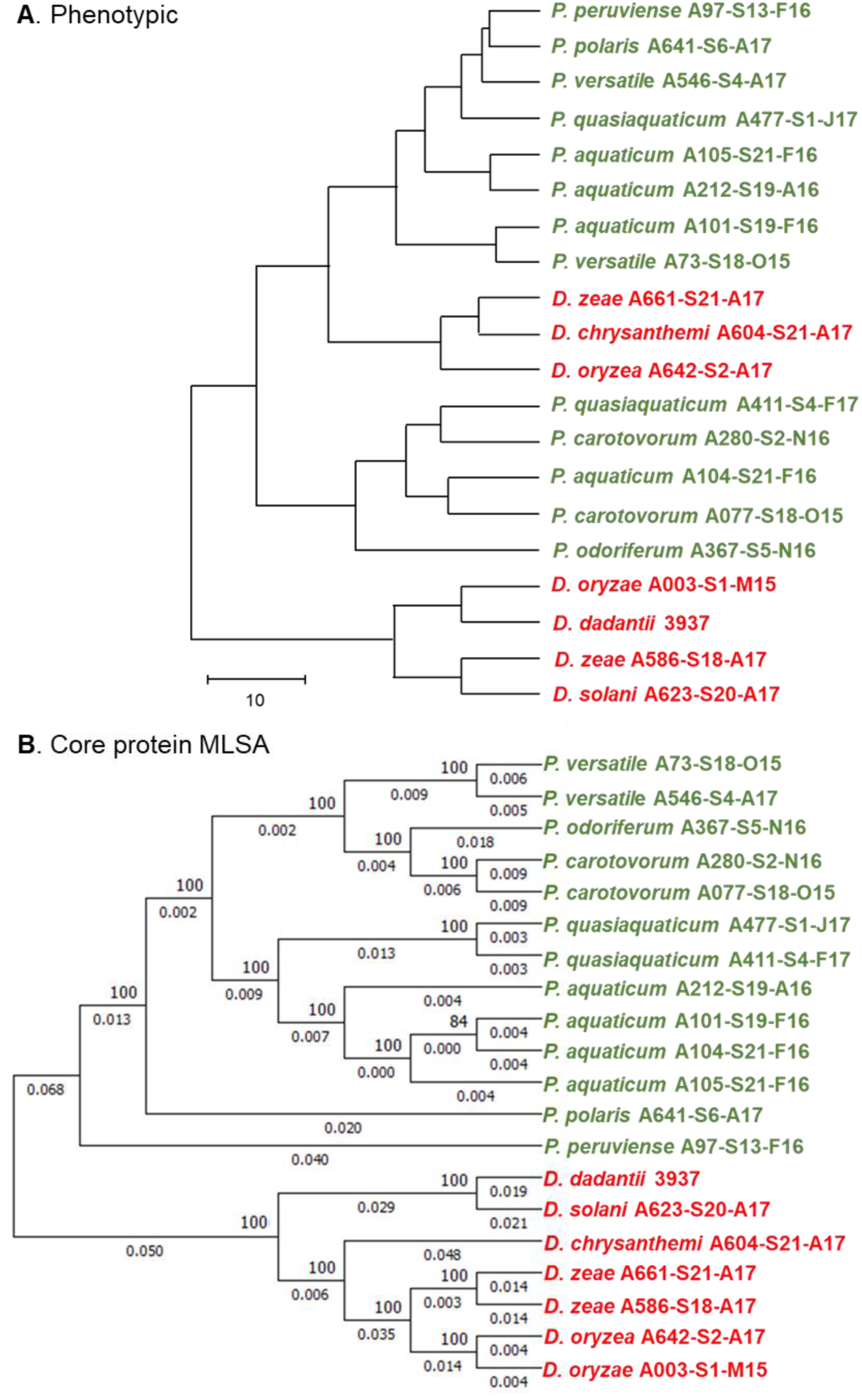
Comparison of phenotypic and phylogenetic trees. (A) dendrogram calculated based on the similarity of killing spectra of investigated strain; the tree is drawn to scale, with branch lengths measured in the distance; (B) multilocus sequence analysis of core nucleotide coding sequences using the BioNJ algorithm. Branch lengths and bootstrap values are shown. *Pectobacterium* spp. strains are marked in green and *Dickeya* spp. strains are marked in red.

**Supplementary Fig. S2.**
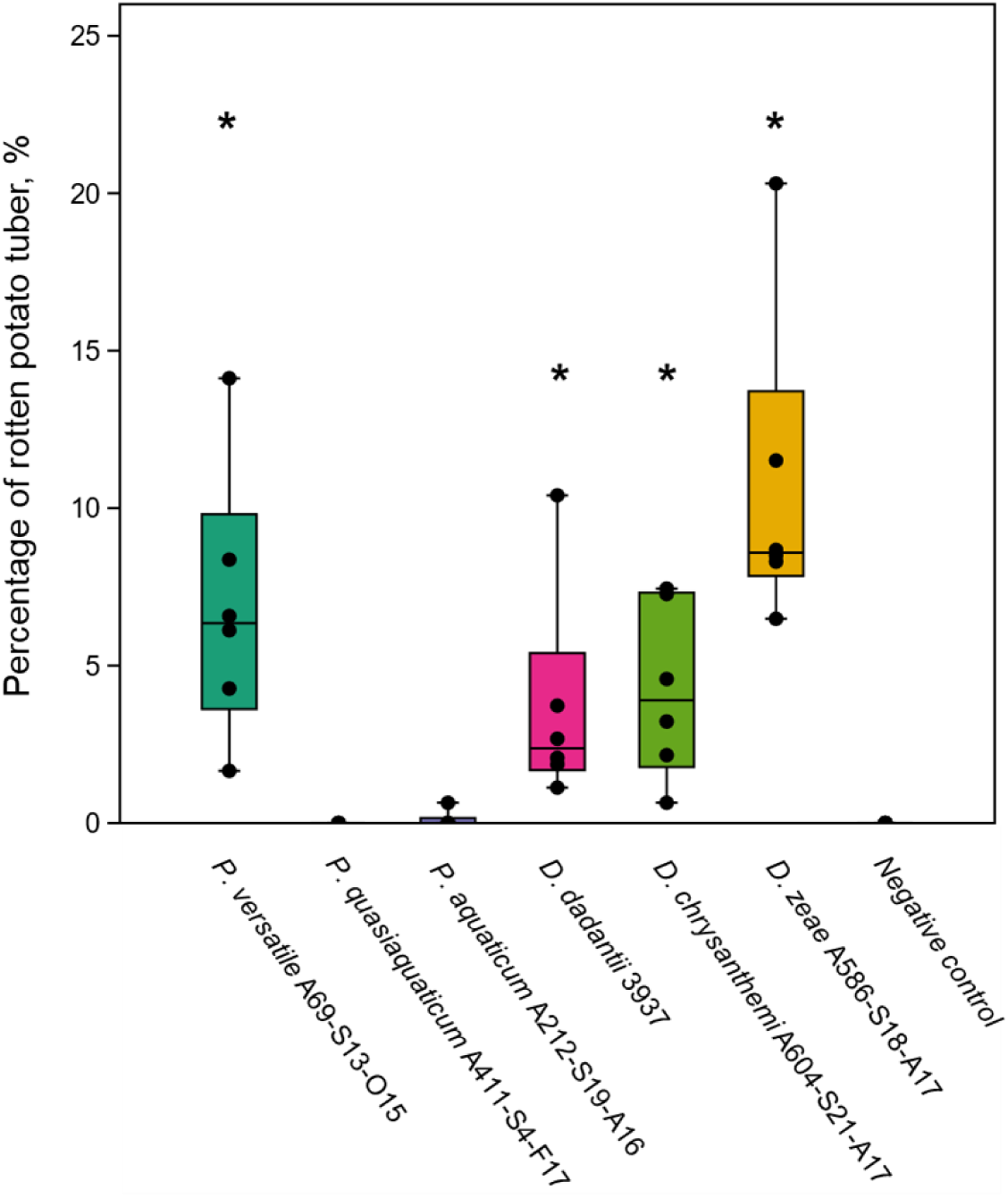
Severity of soft rot symptoms caused on potato tubers by six SRP strains tested for production of tailocins *in planta*. The results are expressed as a percentage of rotten tissue relative to the total mass of each potato tuber. In the box plot, whiskers reflect the maximum and minimum values; the box shows the interquartile range (Q1 to Q3), and the bars inside the boxes represent medians. Points indicate results for individual samples/tubers (n=6). The asterisks (*) denote statistically significant differences (p < 0.05) between a sample inoculated with the given strain and the negative control. Significance was calculated using the one-sample t-test for data sets with normal distribution and the Wilcoxon test for data not following a normal distribution. Negative control – tubers inoculated with sterile PBS buffer.

**Supplementary Fig. S3.**
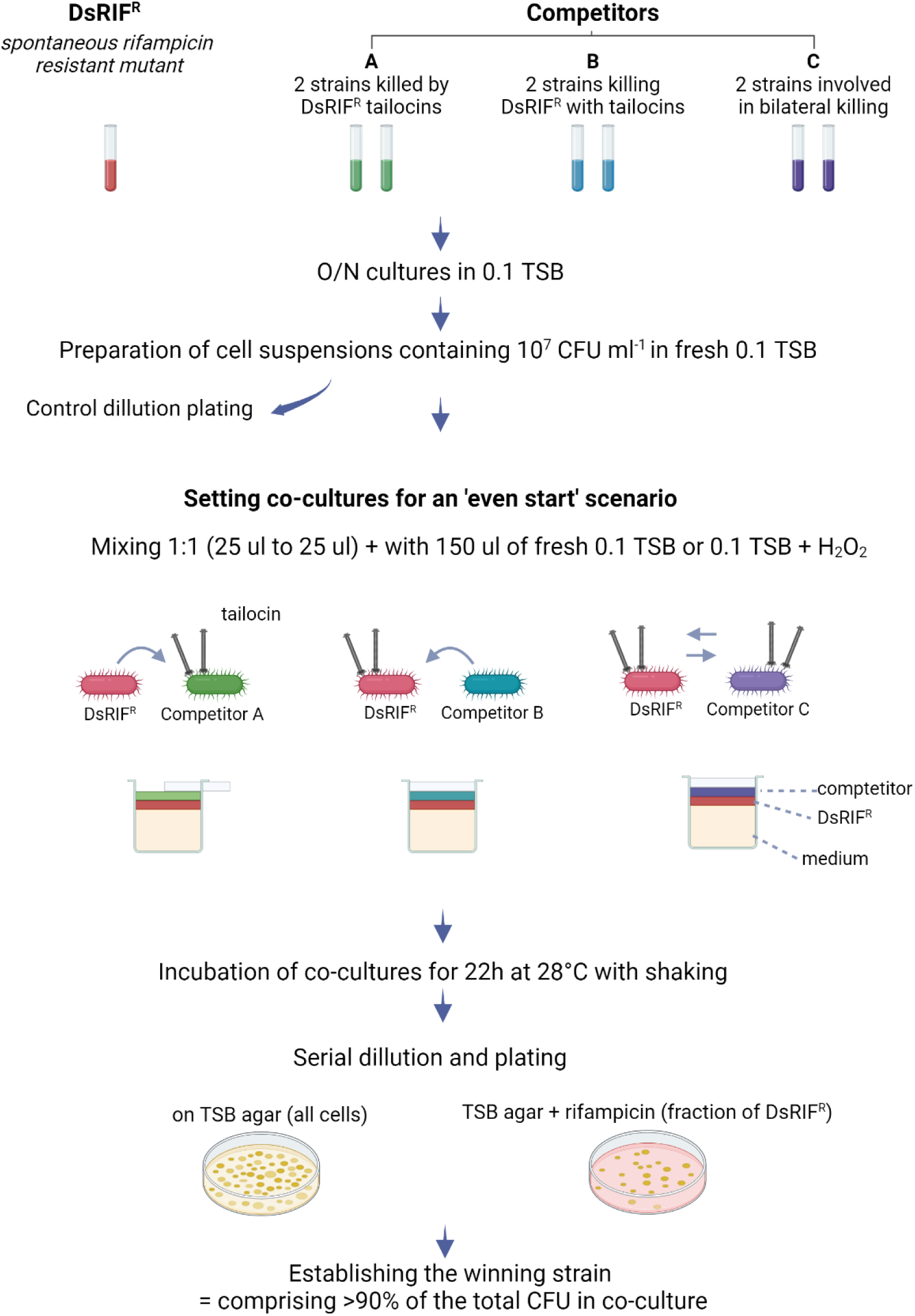
Schematic representation of the steps included in the competition experiment. Created with BioRender.com.

**Supplementary Fig. S4.**
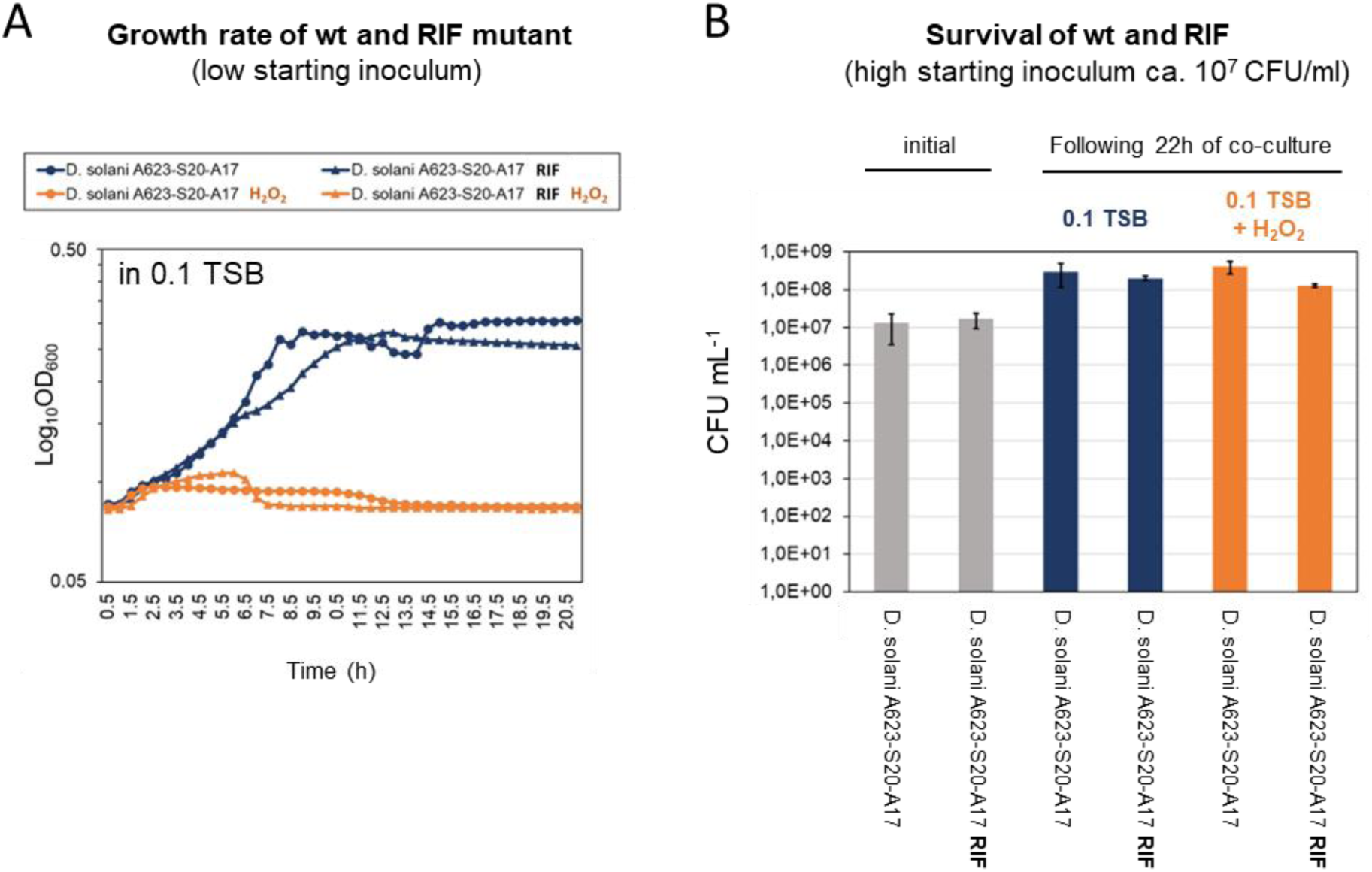
Comparison of growth rates (A) and the survival (B) of the *D. solani* A623-S20-A17 wt and the rifampicin-resistant mutant of this strain in 0.1 TSB alone and supplemented with 0.003 % hydrogen peroxide (H_2_O_2_) The test was performed in 96-well plates incubated at 28°C, with shaking, in a multiwell plate reader (Biotek). To assess growth rates (A), 150 µl aliquots of 0.1 TSB medium were inoculated with 3 µl of suspension containing o/n culture diluted to 0.5 McF (low inoculum). Each time point is an average of two replicates. For cell survival (B), the medium was inoculated with a high initial number of cells (ca. 10^7^ CFU/ml) applied in the co-culture competition experiments (Fig. 5). The results are the average of two separate experiments, 2 technical replicates each. The obtained results indicate that the wt and RIF^R^ variants of *D. solani* A623-S20-A17 have similar growth characteristics in the applied conditions and that 0.003% H_2_O_2_ prevents the growth of the strains when low initial inoculum is used but does not negatively affect the survival of cells in high-density cultures.

## Supplementary datasets

Dataset S1 Results of a megablast search for carotovoricin clusters within 116 closed genomes of *Pectobacterium* spp. The carotovoricin sequence from *P. versatile* A73-S18-O15 was used as a query (genome accession: NZ_CP086369.1).

Dataset S2 Results of a megablast search for dickeyocin clusters within 75 closed *Dickeya* and *Musicola* spp. genomes. Dickeyocin P2D1 from *D. dadantii* 3937 was used as a query (genome accession: NC_014500.1).

Dataset S3 Python script for analyses (ChatGPT-4o)

